# A systematic survey of human tissue-specific gene expression and splicing reveals new opportunities for therapeutic target identification and evaluation

**DOI:** 10.1101/311563

**Authors:** Robert Y. Yang, Jie Quan, Reza Sodaei, Francois Aguet, Ayellet V. Segrè, John A. Allen, Thomas A. Lanz, Veronica Reinhart, Matthew Crawford, Samuel Hasson, GTEx Consortium, Kristin G. Ardlie, Roderic Guigó, Hualin S. Xi

## Abstract

Differences in the expression of genes and their splice isoforms across human tissues are fundamental factors to consider for therapeutic target evaluation. To this end, we conducted a transcriptome-wide survey of tissue-specific gene expression and splicing events in the unprecedented collection of 8527 high-quality RNA-seq samples from the GTEx project, covering 36 human peripheral tissues and 13 brain subregions. We derived a weighted tissue-specificity scoring scheme accounting for the similarity of related tissues and inherent variability across individual samples. We showed that ~50.6% of all annotated human genes show tissue-specific expression, including many low abundance transcripts vastly underestimated by previous array-based expression atlases. As utilities for drug discovery, we demonstrated that tissue-specificity is a highly desirable attribute of validated drug targets and tissue-specificity can be used to prioritize disease-associated genes from genome-wide association studies (GWAS). Using brain striatum-specific gene expression as an example, we provided a template to leverage tissue-specific gene expression to identify novel therapeutic targets. Mining of tissue-specific splicing further reveals new opportunities for tissue-specific targeting. Thus, the high quality transcriptome atlas provided by the GTEx is an invaluable resource for drug discovery and systematic analysis anchored on the human tissue specific gene expression provides a promising avenue to identify novel therapeutic target hypotheses.

## Introduction

Tissue specific genes are those expressed at higher levels in a subset of tissues relative to the baseline expression across all tissues. They often play critical roles in the biological functions unique to those tissues. Identifications of tissue-specific genes have provided a deeper molecular understanding of tissue functions ^1–4^, led to discoveries of key genetic regulatory elements ^5–8^, and defined the molecular basis of human diseases ^9,10^. These studies and on-going gene expression profiling efforts also highlight the usefulness of defining tissue-specific transcriptomic signatures. Therapeutically, pharmacological modulation of tissue-specific genes is a promising avenue to maximize efficacy in target tissues while minimizing the safety risks of affecting unrelated tissues. The availability of a comprehensive catalog of tissue-specific gene expression will set a foundation for a plethora of utilities relevant to human physiology and disease.

Previous efforts to develop a comprehensive transcriptome atlas across multiple human tissue types relied heavily on microarray platforms of limited sample size ^11^. Recent advances in next-generation sequencing technology have enabled a deeper interrogation of the human transcriptome using RNA-seq, with greatly improved sensitivity, accuracy, and a broad dynamic range ^12,13^. Additionally, RNA-seq technology has a unique advantage of allowing transcriptome-wide quantitative assessments of alternative splicing events. The Genotype Tissue Expression (GTEx) project ^14^, an ongoing consortium project, has become the newest flagship of transcriptome atlases by generating the largest cohort of RNA-seq profiles on a broad range of human postmortem tissues. Here, we conducted a comprehensive transcriptome-wide survey to uncover tissue-specific and brain subregion-specific gene expression profiles and alternative splicing events from the GTEx v6 data release, covering 36 human peripheral tissues and 13 brain subregions. Besides a comprehensive characterization of tissue-specific gene expression patterns, we investigated the tissue-specific properties of known drug targets and further focused our analyses on demonstrating the utility of GTEx transcriptome atlas in therapeutic target identification and evaluation. By integrating with GWAS signals for human diseases, we also aimed to identify tissue-specific candidate genes associated with human diseases.

## Results

### Genome-wide quantification of tissue-specificity and brain subregion-specificity from the GTEx human transcriptome atlas

We first sought to derive a comprehensive catalog of human tissue-specific gene expression using the gene-level expression quantification data from GTEx v6p release. After QC and filtering (see Methods), a total of 8527 samples covering 36 peripheral tissues and 13 brain subregions were used for the tissue-specificity analysis. Hierarchical clustering of tissues by their overall gene expression patterns revealed multiple clusters of functionally related tissues with highly correlated gene expression (Figure 1A). In particular, brain sub-regions were highly correlated with each other (correlation coefficient>0.8) compared to other tissues (Figure 1A). With a relatively stringent clustering threshold (see Methods for details), 49 tissues could be clustered into 30 tissue groups (Figure 1A).

**Figure 1.**
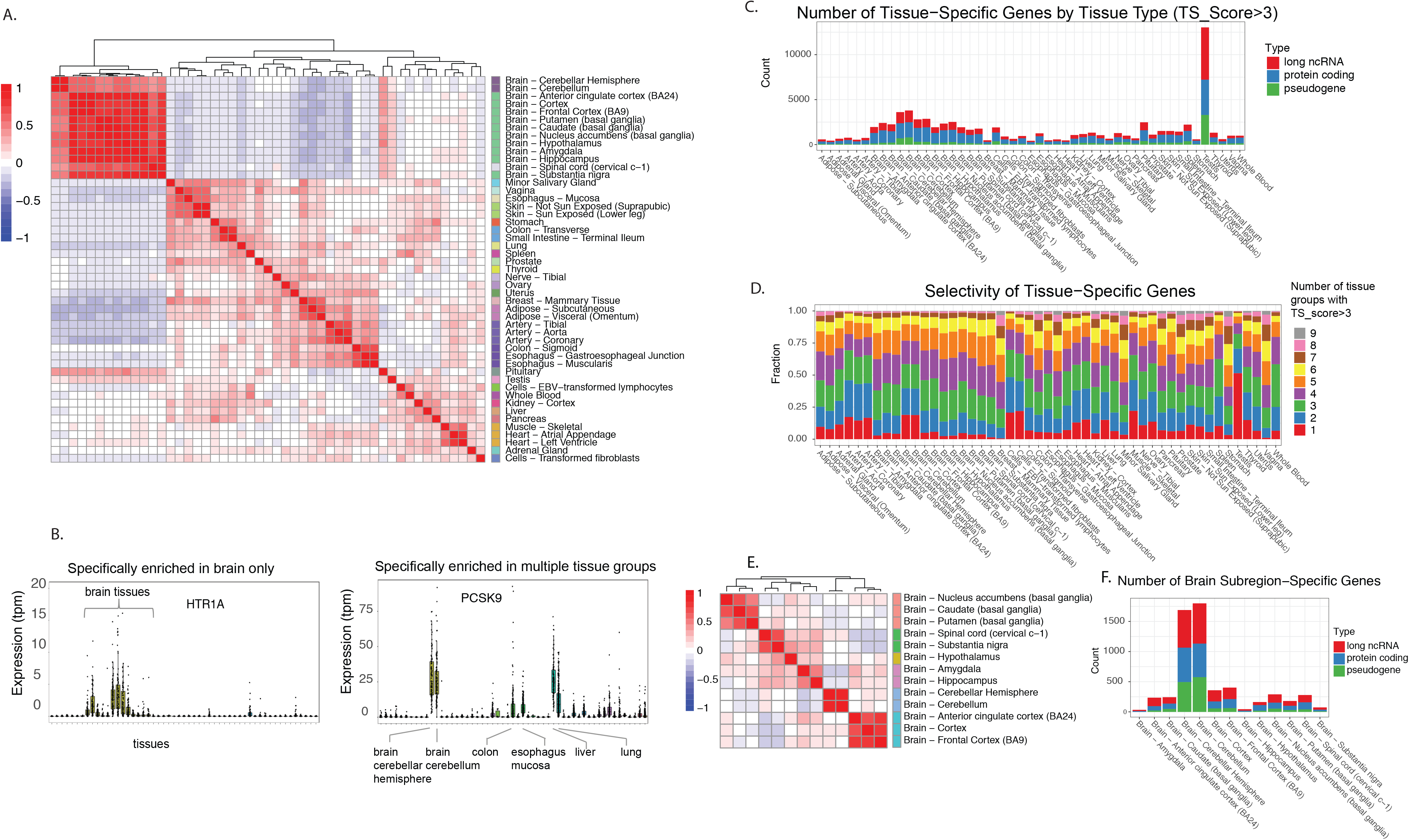
Identification of tissue-specific genes across 49 human tissue types. (A) Clustering of tissues based on overall gene expression specificity patterns. Tissues in the same tissue groups, marked with the color-coded sidebar, show highly correlated global expression pattern. (B) Examples to illustrate the tissue-specific genes identified using a weighted tissue-specificity score. The tissue-specificity scoring method identifies genes specifically enriched in one tissue group as well as genes specifically enriched in a subset of tissue groups. The left panel shows the expression profile of HTR1A, an example gene specifically expressed only in brain. The right panel shows the expression profile of PCSK9, an example gene specifically enriched in a number of tissues. (C) Number of tissue-specific genes identified in each tissue, color-coded by gene type. Tissue-specific genes are defined as TS score>3. (D) Proportions of tissue-specific genes identified in each tissue color-coded by the tissue specificity, defined as the number of tissue groups with TS_Score>3. (E) Clustering of brain subregions based on global gene expression specificity patterns. Color-coded sidebar indicates subregions clustered together (F) Number of subregion-specific genes identified in each brain subregion, color-coded by the gene type.

Several tissue-specificity metrics have been previously reported for microarray-based data ^15–17^. GTEx data, unprecedented in size and nature, required a customized tissue-specificity formula that accounts for transcriptome-wide tissue similarities, donor-to-donor variations, and RNA-seq read count-based statistics. Here we define tissue-specific genes as those showing enriched expression in a subset of tissues compared to typical expression in other tissues. In contrast, we refer to genes present only in a single tissue type as tissue-exclusive genes. Figure 1B illustrates two examples of tissue-specific genes, HTR1A, an example of genes specifically expressed only in brain, and PCSK9, an example of genes specifically enriched in a number of tissues. The mathematical quantification of the tissue-specificity score (TS_Score) is described in detail in the Methods section. Briefly, TS_Score contains three components: relative expression level in one tissue compared to other tissues, tissue-similarity weighting factor, and statistical significance for expression comparison between tissues. TS_Score associated with a gene for a target tissue can be simply interpreted as the average log ratio of expressions in a target tissue over other tissues across all donors. We define tissue-specific genes as having TS_Score greater than 3. We chose the TS_Score>3 threshold as it can be intuitively interpreted as having 8 times (2^3^) the expression in one tissue over the weighted average of the rest of tissues. Statistically, the TS_Score>3 threshold is more than two standard deviations above the mean (μ=−0.05, σ=1.28, see **Supplementary Figure 1** for the overall TS_Score distribution). For the downstream analyses of tissue-specific genes, we further verified our findings could be consistently observed even with a less stringent threshold. To account for the expression similarity among related tissues, we weighted the contribution of each tissue to the tissue-specificity score by how “similar” it is correlated with other tissues. This weighing scheme is particularly important for identifying brain-specific genes due to the overall highly correlated expression patterns across brain sub-regions. Finally, our TS_Score incorporates cross-donor variances to only include differential gene expressions that are statistically significant between the enriched tissue versus all other tissues. This avoids overestimating the tissue-specificity for genes with high variance, especially those expressed at a very low abundance level.

Overall, 50.6% of all protein-coding genes, long noncoding RNA genes, and pseudogenes in the human genome (23569 out of 46508 genes in the GENCODE v19 gene annotation) have TS_Score greater than 3 in at least one of 49 tissues (See the gene expression heatmap of TS genes in **Supplementary Figure 1**). However, the distribution of these genes varies greatly among different tissues (Figure 1C, **Supplemenary Table 1**). Testis had by far the largest number of tissue-specific genes (28% of all genes) with 3923 protein coding genes, 5804 lncRNAs and 3299 pseudogenes. Brain regions also had a higher than average number of tissue-specific genes (11% of all genes; collectively 2855 protein-coding genes, 2367 lncRNAs, and 1225 pseudogenes) most likely reflecting specialized functions of the brain. On the opposite end of the spectrum, adipose, breast, heart, esophagus muscularis, and uterus tissues had the lowest number of tissue-specific genes (<300 protein-coding genes). At the individual gene level, the majority (89.7%) of tissue-specific genes were enriched in less than 5 tissue groups (Figure 1D). Furthermore, only a small number of genes were exclusively enriched in a single tissue type with the highest number in testis.

To verify the functional significance of these tissue-specific genes, we applied functional enrichment analysis on the tissue-specific genes (TS_Score>3) in each tissue type against the Gene-Ontology (GO) annotation. As expected, biological pathways and molecular functions relevant to the tissue were highly enriched in their corresponding tissue-specific genes. For example, genes annotated for “lipid metabolic process” are 5.8-fold enriched in liver-specific genes (pvalue=1.3E-38) and genes annotated for “synaptic signaling” are 5.9-fold enriched in brain cortex-specific genes (Pvalue=3.4E-77).

Given the large number of uniquely identified brain-specific genes, we further interrogated the 13 brain subregions to investigate spatial gene expression relationships. Globally, distinct gene expression patterns of individual brain subregions were observed among interrelated anatomical locations. For example, distinct gene expression clusters were observed among the three functional substructures of the basal ganglia (caudate, putamen and nucleus accumbens), for several areas of cerebral cortex and also cerebellar subregions (Figure 1E). One interesting exception was the observation that genes expressed in the substantia nigra, a small mid-brain region, shared relative high correlation with spinal cord. Surprisingly, among all the brain subregions, the substantia nigra and spinal cord also show higher gene expression correlation with whole blood (**Supplementary Figure 2**), suggesting higher proportion of immune cell types in these regions.

We then further categorized genes with subregion-specific profiles in all 13 brain regions by applying tissue-specificity calculation to only these regions. The cerebellum had the highest number of specifically expressed genes when compared to the rest of the brain (609 protein coding genes and 707 long noncoding RNA genes with TS_Score >3 in cerebellum) with 97% of these genes exclusively enriched in cerebellum (Figure 1E,F, **Supplementary Table 2**). In contrast, less than 50 protein-coding genes with TS_Score >3 were identified in amygdala, hippocampus and substantia nigra, indicating the number of specifically expressed genes in these regions are low and there are similar molecular features among these regions and the rest of the brain. For striatal subregions, hypothalamus, cerebral cortex, and spinal cord, a moderate number of tissue-specific genes (ranging from 50 to 300) were identified.

Annotations of brain subregion-specific genes with biological processes and molecular functions derived from Gene Ontology suggested unique functional enrichment in each of the subregions. For example, cerebral cortex-specific genes were enriched for “synaptic transmission” and “transmission of nerve impulse” including many ion channels and membrane receptors known to regulate neurotransmission. In contrast, genes specifically expressed in the hypothalamus were enriched for “hormone activity”, including neuropeptide and receptor genes (e.g PMCH, HCRT, GHRH, TRH) known to play critical roles in neuroendocrine regulation. Interestingly, genes annotated as “Sequence-specific DNA binding” are also significantly and uniquely enriched in the hypothalamus (enrichment Pvalue=1.3E-5). 12 out of 92 hypothalamus-specific protein-coding genes (e.g. SIX6, SIM1, OTP, HMX2, HMX3) are transcription regulators, suggesting distinct transcriptional regulation in this specialized brain region.

### GTEx transcriptome atlas greatly expands the catalog of tissue-specific gene expression

GTEx, owing to its large sample size and deep sequencing coverage, presents an unprecedented opportunity to discover and annotate tissue-specific genes. Prior to the GTEx project, large-scale gene expression atlases were primarily generated using microarray technology. For comparison, we applied the TS_Score calculation on a highly cited microarray-based atlas, the Genomics Institute of the Novartis Research Foundation (GNF) dataset ^11^. A large number of tissue specific coding (2516) and noncoding genes (10770) were found to be unique in the GTEx dataset as they were not previously measured in the GNF atlas due to the lack of corresponding probes on the microarray platform. In addition, for the 14,891 protein coding genes interrogated in both datasets, more tissue-specific genes were consistently quantified from GTEx as compared to the GNF dataset across all matched tissues. For most tissues, less than half of the tissue-specific genes identified from the GTEx data showed specificity in the GNF atlas (examples shown in Figure 2A). In a few tissues including brain cortex, pancreas, kidney, and testis, greater than 75% of tissue specific genes were found only in the GTEx dataset. In contrast, for most tissues, >90% of tissue-specific genes identified from the GNF atlas were also found to be specific in using the GTEx data. It is unlikely that the GTEx-only tissue-specific genes are due to anatomical isolation differences as isolation or dissection differences would have resulted in the existence of comparable GNF-only tissue-specific genes, which was not observed.

**Figure 2.**
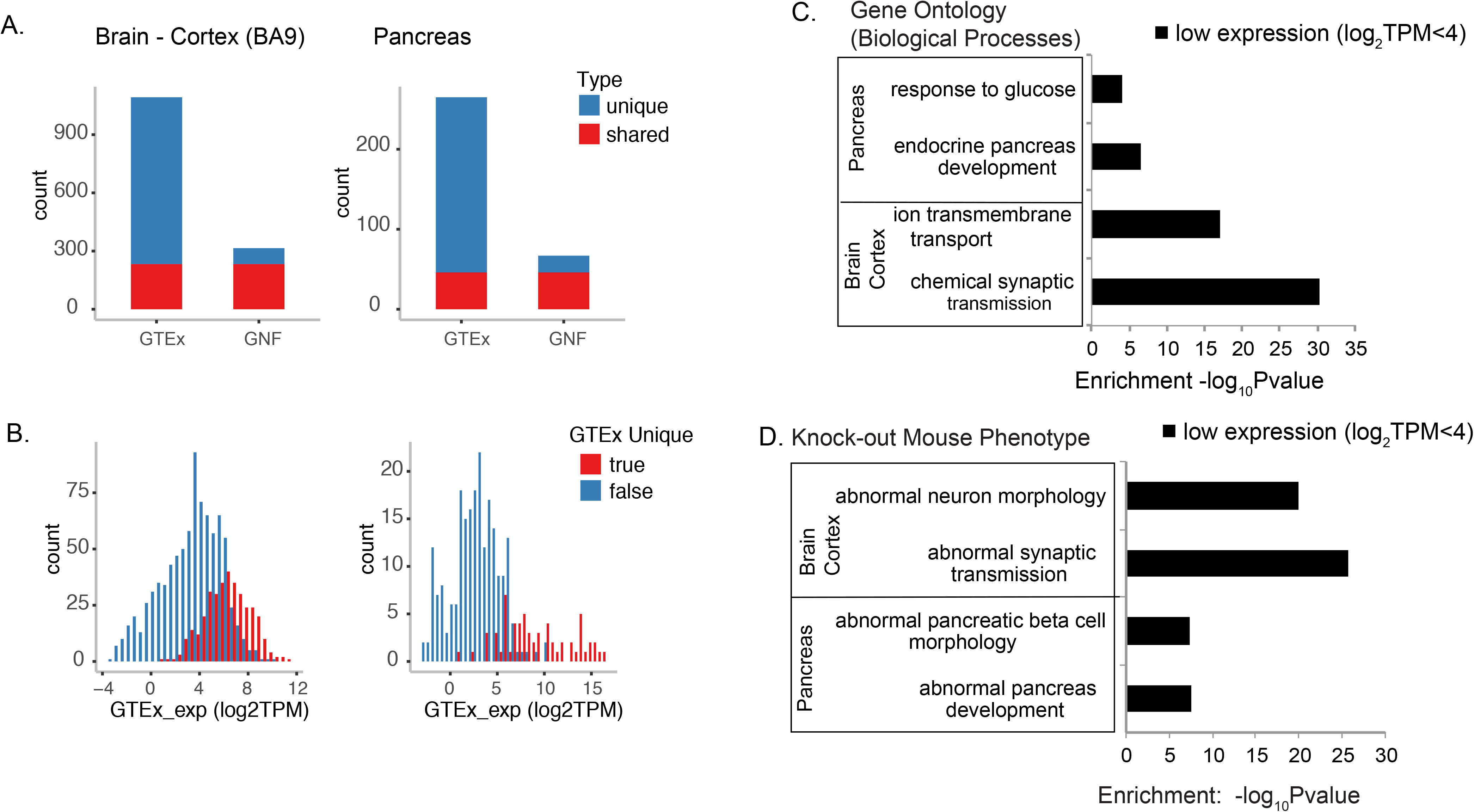
Comparison of tissue-specific genes identified from RNA-seq based GTEx transcriptome atlas with GNF microarray-based tissue expression atlas. Tissue-specific genes identified using GTEx dataset for pancreas, and brain cortex (BA9) were compared with those derived from GNF in the matching tissue types. (A) Overlap of the tissue-specific genes identified from the two datasets showing the large number of tissue-specific genes uniquely identified in the GTEx dataset. (B) Tissue-specific genes uniquely identified in the GTEx dataset have lower abundance than those identified by both datasets. (C) GO functional enrichment of low abundance (log2TPM<4) tissue-specific genes identified in GTEx dataset. (D) Knockout mouse phenotypes enriched in low abundance (log2TPM<4) tissue-specific genes identified in GTEx dataset.

The GTEx-only tissue-specific genes showed a significant shift towards lower expression in all matched tissues (Figure 2B). In fact, the majority (53% on average across tissues) of tissue-specific genes identified in the GTEx data are in the mid-low abundance level (log_2_TPM<4) (Figure 2C), further demonstrating the superior sensitivity and expanded signal dynamic range of RNA-seq. The identification of these numerous low-abundance tissue-specific genes provides fundamental biological insights, as these previously undetected tissue-specific genes may be involved in critical functions within their respective tissues. For example, as indicated by GO enrichment analyses (Figure 2C), “chemical synaptic transmission” is highly associated with low-abundance brain-specific genes (Pvalue=4e-31) while low abundance pancreas-specific genes exert functions towards “endocrine pancreas development” (Pvalue=3e-7). Independent to the literature-driven GO annotations, similar functional enrichment is observed when repeating the analyses using the experiment-driven MGI knockout mouse phenotype database, highlighting the transferability of the findings across species (Figure 2D).

### Tissue-specificity as a desirable attribute for therapeutic drug targets

With the expanded catalog of tissue-specific genes, we next sought to interrogate the tissue-specificity properties of therapeutic drug targets and assess whether tissue specific expression could be used as a criterion for therapeutic target discovery.

We first curated a comprehensive list of drug targets in the different pre-clinical and clinical phases based on its most advanced therapeutic program as documented in the Citeline pharmaproject module. When cross-referenced with our tissue-specificity scores (TS_Score), 68% of the PhIII+ targets (included “PhIII”, “Registered”, “Pre-registration”, “Launched”) were tissue-specific (TS_Score >3) comparing to 46% in all annotated protein coding genes, making PhIII targets 50% more likely to be tissue-specific and a significant enrichment (odds ratio = 2.5, Pvalue=3e-23, Figure 3A, **Supplementary Table 3**). Conversely, we confirmed the reciprocal enrichment that PhIII+ targets were recovered in significantly higher percentage of transcriptome-wide tissue-specific genes comparing to that of the non-tissue specific genes (**Supplementary Figure 4**). Interestingly, cross-sectional survey of targets in earlier drug development stages revealed a significant descend in enrichment for tissue-specificity comparing to PhIII+ targets (Figure 3B).

**Figure 3.**
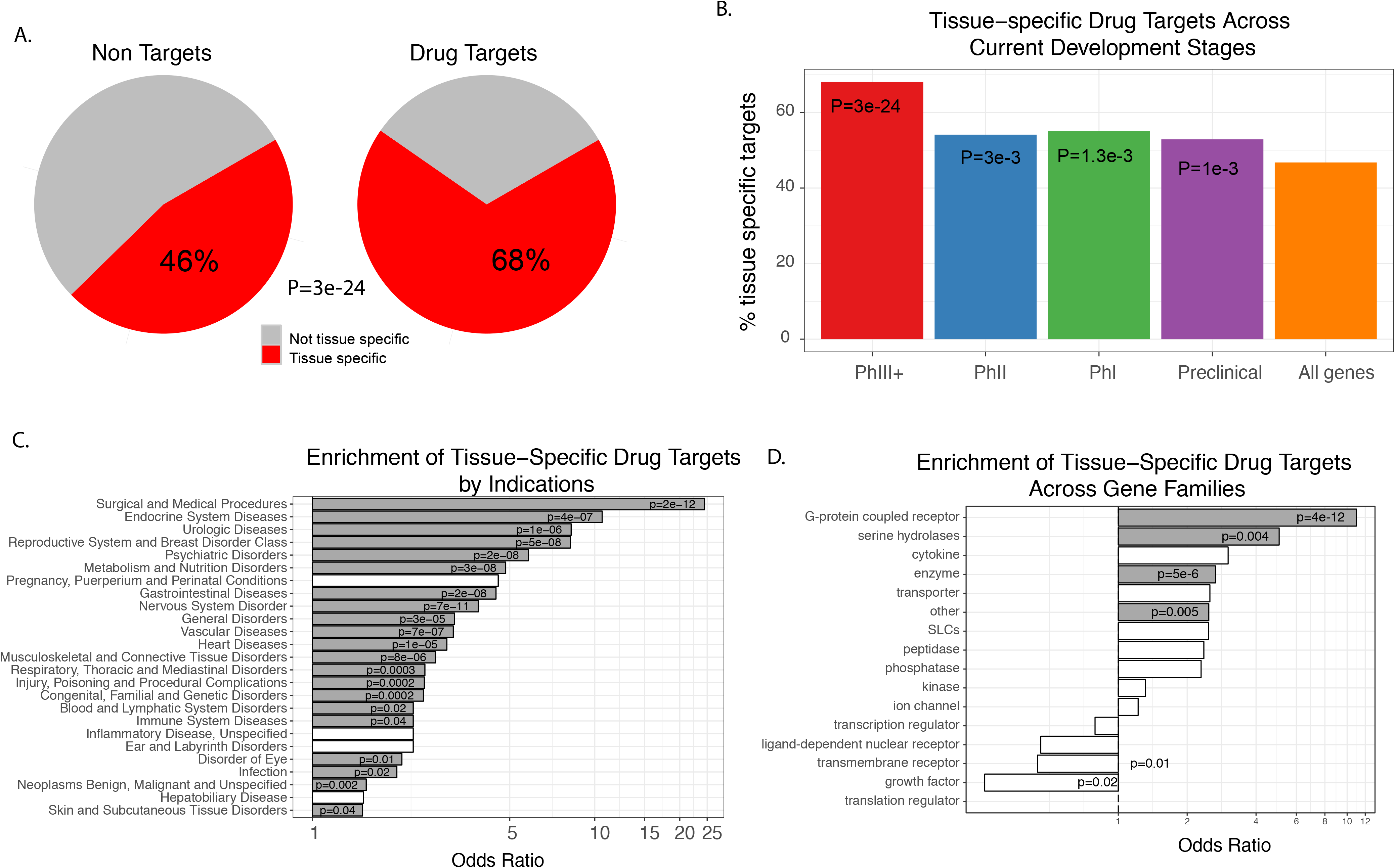
Tissue-specificity and targets of PhIII+ drugs. A) 68% of the PhIII+ drug targets are tissue-specific, significantly more than 46% in the background. B) Targets currently in industry programs before PhIII+ are less enriched for high tissue-specificity (pvalues shown denotes to comparisons with all genes). All phases show significant tissue-specific enrichment comparing to background. Enrichment tissue-specific genes in drug targets of C) different disease indications and D) target class families.

We next investigated which “druggable” gene families were intrinsically tissue-specific by nature. Gene families of the entire GTEx coding transcriptome were annotated according to Ingenuity IPA software. We calculated the median TS_Score based on all members from each gene family. We noted that most of the chemical tractability-aspired and empirically derived “druggable” classes ^18,19^ coincide with high tissue-specificity: GPCRs (median TS_Score=3.6), nuclear receptors (median TS_Score=3.5), ion channels (median TS_Score=4.3), kinases (median TS_Score=2.5), SLCs (median TS_Score=3.5). Further comparisons of the TS_Scores of PhIII+ targets to background within each gene family revealed the preference for even higher tissue specificity for several already highly tissue-specific gene families, including GPCRs (Fisher’s exact adj.p=4E-12), and serine hydrolases (adj.p=0.004) (Figure 3C). To show that this observation was not dependent on the choice of specificity thresholds, we also performed rank-based tests (Mann-Whitney) to test for a shift in TS_Score distribution within gene families. This analysis confirmed the findings and showed a universal trend of upward TS_score shift towards specificity within the PhIII+ drug targets overall (**Supplementary Figure 4C**) and across families (**Supplementary Table 4**).

We postulated that different classes of therapeutic indications likely would have different requirements for tissue-specificity. Figure 3D showed the level of enrichment for tissue-specificity across classes of indications. Of particular note, endocrine system diseases, driven by hormone-impaired indications, showed that PhIII+ targets were 10 times more likely to have a strong preference for tissue-specificity through targeting the hormone receptors (GHRHR, PTH1R, SSTR family, etc). Psychiatric targets, as expected, were highly enriched for tissue-specificity (OR>5), and were dominated by brain-specific genes. Interestingly, Metabolic indications represent one of the larger collections of PhIII targets (spanning 147 distinct targets), which collectively showed preference for high tissue-specificity (0R>4) concentrating in artery, heart, adipose, and lymphocytes. In contrast, indications in skin and oncology showed the least preference for tissue-specific targets, perhaps owing to local delivery mechanism and lower toxicity hurdle, respectively.

### Tissue-specificity of druggable target gene families and utilities in target identification

Historically, small molecule drug discovery efforts have been focusing on several major target gene families including G-protein coupled receptors (GPCRs), kinases, metabolic enzymes such as serine hydrolases (SHs), solute transporters (SLCs), and ion channels, as they are more amenable for small molecule intervention ^20,21^. Interestingly, by overlaying the tissue specificity profiles with these druggable gene families, we observed distinct tissue-specific distribution patterns of the different drug target classes (Figure 4A, **Supplementary Table 5**). Brain tissues have the largest number of specifically expressed ion channels, GPCRs (best enrichment Pvalue =1.4e-16), ion channels (best enrichment Pvalue =2.8e-56, solute transporters (best enrichment Pvalue =1.1e-4), many of which are involved in neuronal signal transmission processes. In contrast, tissue-specific serine hydrolases are highly enriched in pancreas (Pvalue=7.4e-11), vagina (Pvalue=2.3e-10), liver (Pvalue=5.8e-10), skin (Pvalue=1.6e-9), and esophagus (Pvalue=5.6e-9), but not in brain. For peripheral tissue types, tissue specific ion channels are also highly enriched in testis (Pvalue=1e-7), skeleton muscle (9.8e-7), heart (Pvalue=4.1-6), colon (3e-6), tissue-specifc GPCRs highly enriched in spleen (Pvalue=6.9e-15), whole blood (Pvalue=2.2e-9) and tissue-specific solute transporters highly enriched in kidney (Pvalue=5.6e-23), liver (Pvalue=1.8e-11), small intestine (Pvalue=4.3e-8). In contrast, tissue-specific kinases are not significantly enriched in any central or peripheral tissue types. These distinct tissue specificity patterns reflect the specialized functions of the tissues, and could drive novel therapeutic target identification.

**Figure 4.**
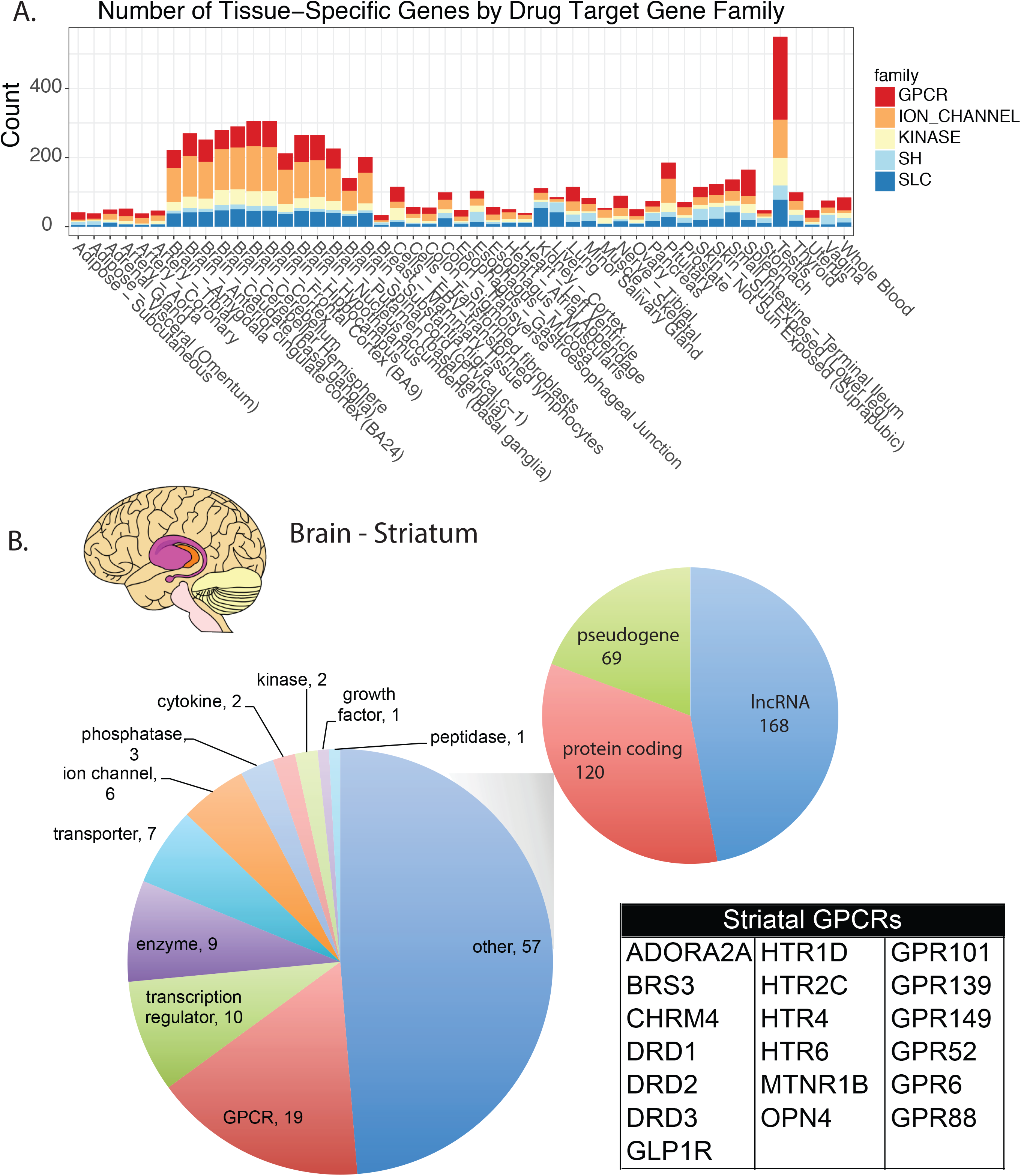
Identification of tissue-specific genes in drug target classes for potential therapeutic targets. (A) Number of tissue-specific genes in different drug target gene families. (B) A focused view of striatum-specific genes identified. Pie charts show the breakdown of striatum-specific genes by gene type and major gene family. The gene table on the right shows the large number of striatum-specific GPCR genes identified including several orphan GPCRs.

Here, we took a closer examination of the tissue-specific GPCRs as an illustration of a tissue-specificity driven approach for target identification. 312 (85%) out of a total of 369 non-olfactory GPCRs are tissue-specifically expressed (1.8 fold over the overall 46.8% tissue-specific genes in protein-coding genes) with the largest number of them and highest enrichment found in brain regions and immune systems, i.e. whole blood, lymphocytes, spleen (enrichment Pvalue<2e-9) Of these, 80 are annotated as orphan GPCRs (oGPCRs). The endogenous ligands for many of the oGPCRs are elusive and their biological functions are largely unknown. Yet, ~27% of all FDA-approved drugs are targeting only ~50 GPCRs with known ligands. Thus, oGPCRs represent many untapped opportunities for novel therapeutic targets. These tissue-specific oGPCRs are particularly interesting, as their tissue-specific expression pattern provides additional context to elucidate their biological function and hypothesize their therapeutic potentials. An interesting case example can be derived from the striatum of the brain. The striatum is an evolutionarily conserved subcortical structure that performs important functions including movement control, regulation of attention, motivation and cognition, and the processing of rewarding and salient stimuli ^22^. Disruption of striatal function is associated with many neurological and psychiatric diseases including Parkinson’s and Huntington’s disease, addiction, and psychosis ^23^. A deeper gene expression analysis on the three substructures of human striatum (caudate, putamen and nucleus accumbens) identified a total of 120 protein-coding genes, 168 long non-coding RNAs and 69 pseudogenes specifically expressed in the striatum (Figure 4B). Functional annotation of striatal specific protein-coding genes showed strong enrichment in dopamine neurotransmission and signaling (e.g. DRD1, DRD2, DRD3, PDE1B, PDE10A, PPP1R1B, RGS14), consistent with the well-described function of the striatum to coordinate dopamine-dependent brain functions. Strikingly, a large number of GPCRs are highly specifically expressed in the human striatum (Figure 4B). Out of the 120 protein coding genes identified as specifically expressed in striatum, 19 of them are GPCRs. Out of these GPCRs, 13 have known ligands including many established therapeutic or investigational drug targets (e.g. dopamine D1, D2, D3 receptors, serotonin 5-HT2C, 5-HT1D, 5-HT6 receptors, and others). Interestingly, 6 of the remaining GPCRs are all orphan receptors (**Supplementary Figure 5**), namely, GPR6, GPR52, GPR88, GPR101, GPR139 and GPR149 whose endogenous ligands and functions are unknown. Given that many of the 13 striatum-specific GPCRs with known ligands play key roles in modulating striatal neurotransmission and striatal functions, the identification of these 6 orphan GPCRs provides a new set of potential drug targets whose modulation may be therapeutic for treating striatal-related neurological diseases.

### Overlap of GWAS candidate genes associated with human diseases with tissue-specific genes

Recent progresses in GWAS studies of human diseases have enabled more human genetics driven therapeutic target discovery ^24,25^. Here we further examined the tissue-specificity profile of genes near the GWAS signals and determined whether GWAS candidate genes for human diseases are enriched in tissue-specific genes. We first curated the GWAS findings for 61 diverse human diseases, disease-associated biomarkers, or human traits from the NHGRI catalog (See Method for the curation details). As a first approximation, for each GWAS locus, we assigned a single candidate gene as the one that is closest to the GWAS lead SNP (i.e. Single Nucleotide Polymorphism with the most significant GWAS Pvalue). Then for each tissue type, we assessed the overlap of tissue-specific genes with the GWAS candidate genes for each of the 61 diseases/traits. Remarkably, GWAS signals are often significantly enriched in tissue-specific genes in the tissue types that are directly relevant to the disease (Figure 5, Table 1). This enrichment pattern can be consistently observed even when less stringent TS_Score threshold was used to define tissue-specific genes (**Supplementary Figure 6**). For example, for Systemic Lupus Erythematosus (SLE), an autoimmune disease triggered by immune cells, 6.4-fold enrichment (Pvalue=1.78E-6) of the GWAS candidate genes were observed only in lymphocyte-specific genes. In fact, one third of the GWAS candidate genes (10 of 32) are specifically expressed in lymphocyte. In comparison, for abnormal heart rate, the strongest enrichments (~21-fold, Pvalue=2.62E-6) were observed in genes specifically expressed in heart tissues (e.g. heart atrial appendage). Interestingly, among the candidate genes associated with the levels of various metabolites, homocysteine, urate, cholesterols and triglycerides, most significant enrichments (9-12 folds, Pvalues range 5E-4 to 3E-10) were observed in genes specifically expressed in liver and kidney, the two key organs responsible for the metabolic homeostasis. For Type-1-diabetes (T1D) and Type-2-diabetes (T2D) GWAS signals are distinctly enriched around tissue-specific genes from different tissues. For T1D, the strongest enrichment (4.5 fold, Pvalue=1E-4) was observed in lymphocyte-specific genes. In contrast, for T2D, moderate enrichments (~3.6-fold, Pvalue~0.006) were found in genes specifically expressed in pancreas, but no significant enrichment in whole blood or lymphocyte-specific genes. This matches remarkably well the difference of the etiology of the two diseases. T1D is considered an autoimmune disease where beta cells are attacked by the immune system, while T2D is likely due to dysfunctional metabolism. Surprisingly, no strong enrichment was observed in any tissue type for any obesity related GWAS loci, with only marginally enriched signals among the tissue-specific genes in brain sub-regions.

**Figure 5.**
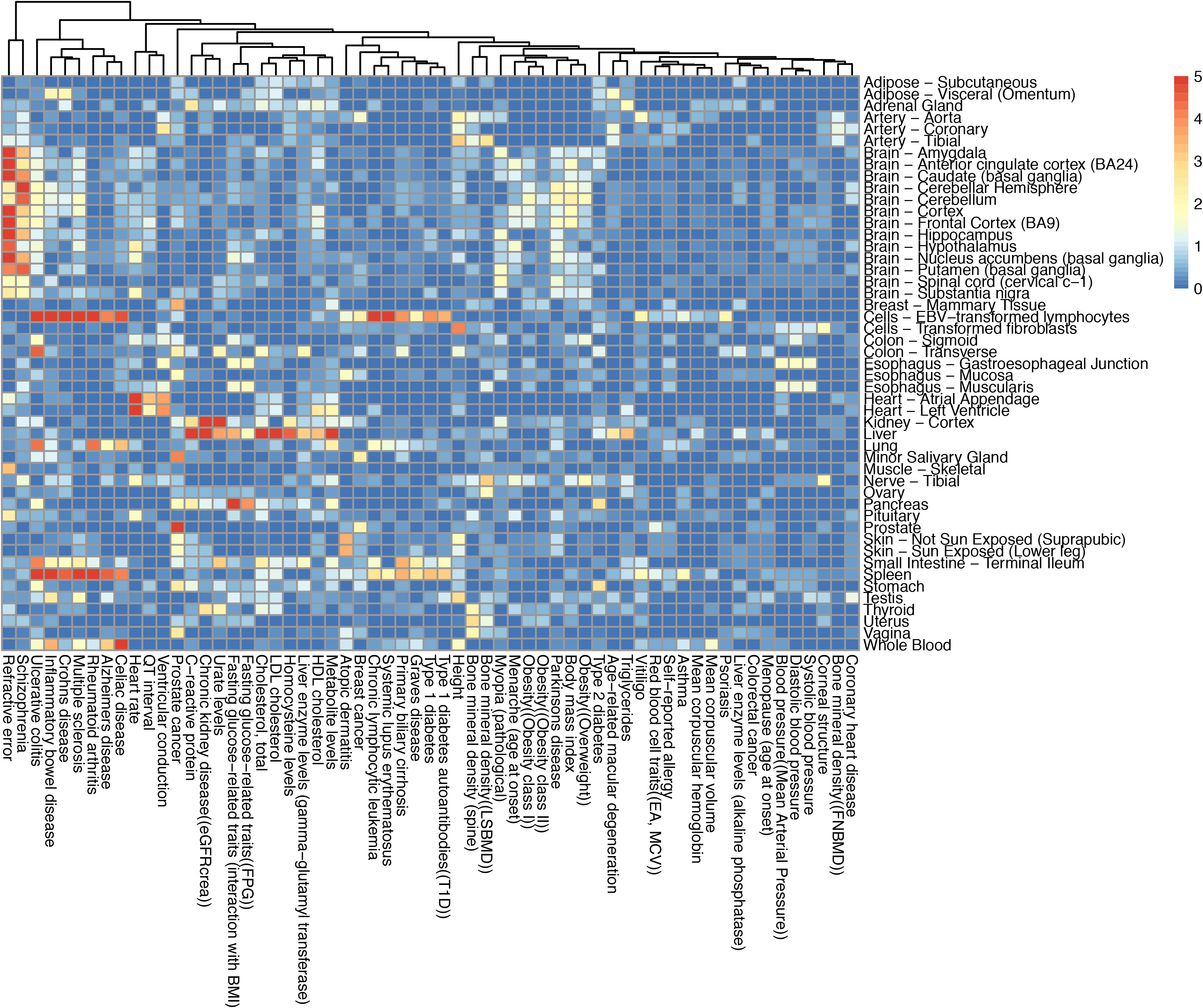
Tissue-specific genes are enriched for GWAS signals for human diseases and traits. Heatmap is colored by the significance (−log_10_Pvalue) of the overlap between tissue-specific genes (TS_Score>3) and the candidate genes closest to GWAS peak signals for 61 human diseases and traits.

**Table 1.** List of tissue-specific genes enriched in the GWAS loci for human diseases and traits. For each human disease and trait that showed significant enrichment of tissue-specific genes in the corresponding GWAS risk loci, the table shows the tissue with the strongest enrichment, the enrichment statistics, and the list of tissue-specific genes located adjacent to the GWAS peak signals.

| Tissue | GWAS Traits/Diseases | # of tissue-specific genes | # of GWAS candidate genes (closest gene to the GWAS peak SNPs) | Overlap | Enrichment P-value | Overlap Genes |
| --- | --- | --- | --- | --- | --- | --- |
| Cells - EBV-transformed lymphocytes | Alzheimers disease | 977 | 23 | 7 | 8.0E-05 | MEF2C,CR1,SORL1,GPR141,INPP5D,PTK2B,HLA-DQA1 |
| Spleen | Alzheimers disease | 1154 | 23 | 8 | 2.7E-05 | SLC24A4,CR1,ABCA7,GPR141,MS4A6A,INPP5D,CD33,HLA-DQA1 |
| Skin - Not Sun Exposed (Suprapubic) | Atopic dermatitis | 872 | 15 | 5 | 3.2E-04 | FLG,OVOL1,CYP24A1,LCE5A,OR10A3 |
| Liver | C-reactive protein | 635 | 18 | 6 | 1.3E-05 | CRP,GCKR,HNF1A,SALL1,HNF4A,APOC1 |
| Cells - EBV-transformed lymphocytes | Celiac disease | 977 | 26 | 8 | 2.2E-05 | PLEK,RGS1,IL12A,C1orf106,HLA-DQA1,ICOSLG,RMI2,RUNX3 |
| Liver | Cholesterol, total | 635 | 50 | 14 | 3.0E-10 | PCSK9,LIPC,HNF4A,LIPG,ABCG8,HNF1A,SLC22A1,HPR,CYP7A1,NPC1L1,APOB,GCKR,NAT2,APOC1 |
| Kidney - Cortex | Chronic kidney disease(eGFRcrea) | 549 | 21 | 7 | 9.4E-07 | SLC22A2,SLC34A1,SLC6A13,UMOD,WDR72,NAT8,SLC7A9 |
| Cells - EBV-transformed lymphocytes | Chronic lymphocytic leukemia | 977 | 16 | 8 | 2.9E-07 | ODF1,SP140,ACOXL,IRF4,BCL2,IRF8,PMAIP1,HLA-DQA1 |
| Cells - EBV-transformed lymphocytes | Crohns disease | 977 | 96 | 17 | 3.6E-06 | IL23R,GPR65,TNFSF11,IL12B,KIF21B,ICOSLG,IL10,HLA-F,ZPBP2,LTA,B3GNT2,SP140,NOD2,RASGRP1,HLA-DQA2,MUC19,IL2RA |
| Liver | Fasting glucose-related traits (interaction with BMI) | 635 | 19 | 5 | 2.6E-04 | GCKR,PDX1,SLC2A2,PROX1,FOXA2 |
| Pancreas | Fasting glucose-related traits (interaction with BMI) | 364 | 19 | 6 | 7.7E-07 | SLC30A8,PDX1,SLC2A2,C2CD4A,G6PC2,FOXA2 |
| Liver | HDL cholesterol | 635 | 45 | 7 | 5.0E-04 | LIPC,HNF4A,LIPG,APOB,F2,APOC1,LPA |
| Heart - Left Ventricle | Heart rate | 259 | 18 | 5 | 2.6E-06 | CHRM2,HCN4,CCDC141,NKX2-5,MYH6 |
| Liver | Homocysteine levels | 635 | 13 | 5 | 3.3E-05 | HNF1A,CPS1,CBS,SLC17A3,FGF21 |
| Cells - EBV-transformed lymphocytes | Inflammatory bowel disease | 977 | 102 | 23 | 5.6E-10 | IL23R,GPR65,IKZF1,IKZF3,IL12B,ICOSLG,PRKCB,CD226,LPXN,MUC19,GPR183,HLA-DRB1,CD40,FEN1,LSP1,IL10,RNASET2,IFNG,IRF8,CXCR5,NOD2,C1orf106,IL2RA |
| Liver | LDL cholesterol | 635 | 36 | 9 | 1.4E-06 | SLC22A1,PCSK9,HPR,CYP7A1,NPC1L1,APOB,ABCG8,HNF1A,APOC1 |
| Liver | Metabolite levels | 635 | 30 | 11 | 9.6E-10 | SERPINA1,TAT,HPR,LIPC,GC,APOB,GCKR,F12,TM4SF5,APOC1,KLKB1 |
| Cells - EBV-transformed lymphocytes | Multiple sclerosis | 977 | 50 | 16 | 9.6E-10 | CD86,GPR65,PLEK,RGS1,IL12B,HLA-F,IRF8,SP140,CLECL1,CD58,C1orf106,HLA-DQA1,TNFSF14,BATF,IL2RA,HLA-DRA |
| Cells - EBV-transformed lymphocytes | Primary biliary cirrhosis | 977 | 24 | 7 | 1.1E-04 | CXCR5,IKZF3,SPIB,POU2AF1,IRF8,IL12RB2,HLA-DQB1 |
| Prostate | Prostate cancer | 471 | 51 | 10 | 2.6E-07 | SALL3,MSMB,NKX3-1,MMP7,HNF1B,KLK3,BIK,IRX4,GRHL1,MLPH |
| Heart - Atrial Appendage | QT interval | 310 | 11 | 3 | 5.5E-04 | KCNH2,SCN5A,SLC8A1 |
| Brain - Cortex | Refractive error | 1545 | 20 | 10 | 6.6E-07 | PTPRR,RBFOX1,RASGRF1,RORB,SHISA6,GJD2,CYP26A1,ZMAT4,KCNQ5,GRIA4 |
| Cells - EBV-transformed lymphocytes | Rheumatoid arthritis | 977 | 34 | 13 | 3.0E-09 | BLK,CD40,CD83,CSF2,PLD4,PTPN22,CCR6,TRAF1,IL2RA,B3GNT2,HLA-DRA,NFKBIE,HLA-DQB1 |
| Brain - Putamen (basal ganglia) | Schizophrenia | 1119 | 109 | 18 | 3.3E-05 | RIMS1,PPP1R16B,DGKI,GRIA1,GPM6A,CACNA1I,GRIN2A,FUT9,DRD2,BCL11B,SNAP91,ZNF536,NRGN,GRM3,C11orf87,CSMD1,ANKRD63,HCN1 |
| Cells - EBV-transformed lymphocytes | Systemic lupus erythematosus | 977 | 32 | 10 | 1.8E-06 | LRRC18,CD80,TNFAIP3,IKZF1,RASGRP3,TNFSF4,IRF5,HLA-DQB2,HLA-DRB1,GPR19 |
| Liver | Triglycerides | 635 | 33 | 6 | 5.3E-04 | LIPC,APOE,APOB,GCKR,CYP26A1,NAT2 |
| Cells - EBV-transformed lymphocytes | Type 1 diabetes | 977 | 41 | 9 | 1.3E-04 | PTPN22,IFIH1,CTSH,HLA-DQA1,IL10,CENPW,HLA-DRA,CD226,CD69 |
| Cells - EBV-transformed lymphocytes | Ulcerative colitis | 977 | 55 | 13 | 1.8E-06 | IL23R,LSP1,ITGAL,IL12B,ICOSLG,IL10,IFNG,ZPBP2,IRF8,IRF5,C1orf106,HLA-DQA1,HLA-DRA |
| Kidney - Cortex | Urate levels | 549 | 28 | 8 | 5.9E-07 | HNF4G,LRP2,SLC17A1,SLC22A11,TMEM171,PDZK1,OVOL1,A1CF |
| Heart - Left Ventricle | Ventricular conduction | 259 | 20 | 4 | 1.1E-04 | TBX5,HAND1,TBX20,CASQ2 |

For neurological disorders, such as Parkinson’s disease and schizophrenia, the strongest enrichments of GWAS signals were observed in brain-specific genes. For schizophrenia in particular, GWAS signals from the recent Psychiatric Genomics Consortium study ^26^ were located near several brain enriched ion channels and receptors (CACNA1I, GRIN2A, HCN1, DRD2, GRM3), and synaptic proteins (SNAP91, NRGN, RGS6, RIMS1) that are known to play important roles of neurotransmission. Perhaps the most surprising enrichment was observed in the GWAS loci for late onset of Alzheimer’s disease (LOAD). Instead of enrichment in brain-specific genes, the strongest signal was observed for genes specifically expressed in the immune system (Table 1). Out of the 23 candidate genes from the AD GWAS loci, 11 of them (ABCA7, PTK2B, CD33, CR1, GPR141, INPP5D, MS4A6A, SLC24A4, SORL1, MEF2C, HLA-DQA1) are highly enriched in whole blood, lymphocyte, and spleen (Pvalue<6E-4), suggesting a causal link between immunity and the development of LOAD.

In total, 33 out of the 61 diseases/traits showed significant enrichment of GWAS signals (Pvalue<0.001) around tissue-specific genes, accounting for 31% of GWAS loci. The full list of tissue-specific GWAS candidate genes is included in Table 1. Taken together, our results strongly indicate that tissue-specific genes identified are highly relevant to human diseases, and tissue-specific gene expression provides an orthogonal criterion for mapping candidate disease genes for GWAS loci to uncover causal links between genes and human diseases, and developing potential therapeutic target hypotheses.

### Tissue-specific splicing provides novel opportunities for tissue-specific targeting

It is well established that majority of the multi-exon genes in the human genome undergo alternative splicing. Translation of the different splice isoforms of the same gene often lead to different protein products that play important biological functions and exert different pharmacological effect in the various tissues and cell types. Prior to GTEx, resources for quantitative and comprehensive assessment of splice isoform distribution in human tissues are limited.

To further assess alternative splicing, we applied a comprehensive workflow with a focus on quantifying all alternative splicing (AS) events including alternative exons and exon skipping (AltEx), alternative 5’ and 3’ splice site (Alt5_3), and intron retention (IR) (see Methods for details, **Supplementary Figure 7**), as opposed to reconstruction and quantification of the full length isoform. AS event-based quantification is more robust against various confounding factors such as RNA degradation and fragment length biases, and is less affected by the incomplete annotation of alternative splice isoforms in the reference annotation ^27^. We first identified and quantified all splicing events in each sample according to their types using the following metrics, PSI: percent spliced in (for AltEx events), PSU: percent splice site usage (for Alt5_3 events), PIR: percent intron retention (IR events) (See Methods for the analysis workflow and **Supplementary Figure 7** for a schematic description of the different types of AS events and the formula). Then for each splicing event, the tissue PSI/PSU/PIR values were calculated as the median PSI/PSU/PIR values across the samples of each tissue type. The total set of AS events across all tissue types are defined as splicing events having tissue PSI/PSU/PIR<90% and >10% in any of the tissue types and/or having a range of tissue PSI/PSU/PIR values >25% across all tissues ^28^. Overall, 74890 AS events (20255 AltEx, 9401 Alt5_3, 39575 IR events, Figure 6A) were detected in 13,274 genes in the 49 tissue types. As previously reported ^29^, we observed widespread intron-retention events across all tissue types, accounting for 60% of all AS events detected. Since GTEx RNAseq libraries were purified using a polyA enrichment method that enriches for mature polyadenylated mRNA and a large number of samples were profiled for each tissue type, we reasoned that these intron-retention events consistently observed across tissue samples were unlikely to be artifacts of pre-mRNA contamination.

**Figure 6.**
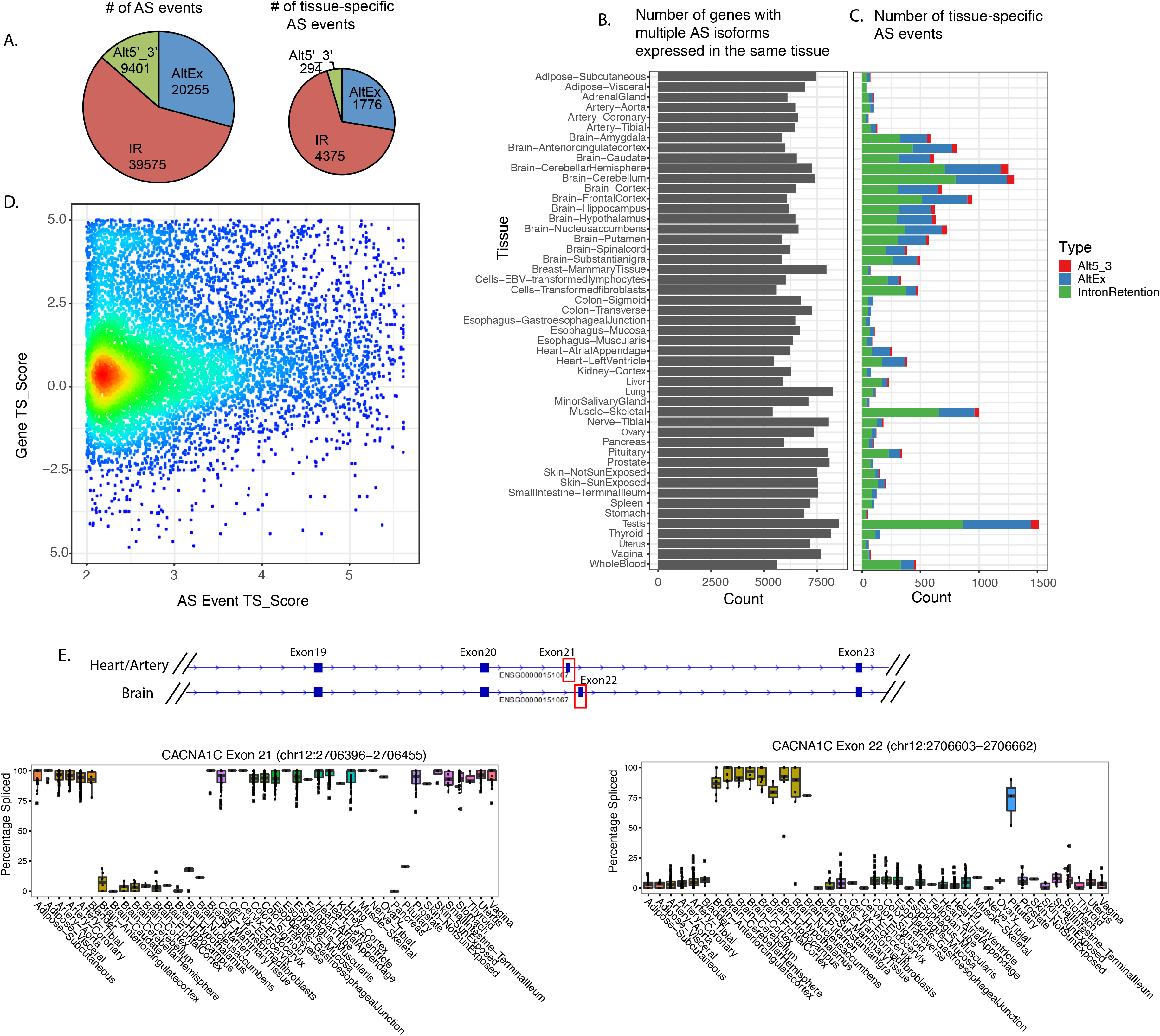
Alternative splicing events across human tissue types, their tissue-specificity and utility for tissue-specific targeting. **A**) Number of different types of AS events and tissue-specific AS events identified across GTEx tissues. AltEx: alternative exon and exon/microexon skipping events, Alt5’_3’: alternative 5’ or 3’ splice site, IR: intron retention B) Number of genes with multiple AS isoforms expressed in same tissue. C) Number of tissue-specific AS events identified in each tissue, color-coded by the type of alternative splicing. D) Distribution of gene-level tissue specificity score (gene TS_Score) for genes with tissue-specific AS events. For clarity, only those with AS event TS_Score>2 were shown. E) Brain specific alternative splicing in CACNA1C exon21 and exon22, as an example of considering alternative splicing in a non-brain-specific gene enables brain-specific targeting of drug targets.

The number of genes with multiple AS isoforms simultaneously expressed in a tissue (i.e. genes having at least one AS event with 10%<PSI/PSU/PIR value<90% in the tissue) is rather comparable across the diverse tissue types (6778 genes on average ± 826 genes, Figure 6B). Despite the complex and higher order functions of the human brain, no substantial increase in the number of genes expressing multiple AS isoforms was observed in any of the brain regions. However, when clustering tissues based on the transcriptome-wide splicing patterns (i.e. using the PSI/PSU/PIR values), brain regions clustered separately from the peripheral tissues, indicating that the splicing patterns in brain regions are distinct from peripheral tissues (**Supplementary Figure 8**). Overall, anatomically related tissues generally show more similar splicing patterns in the clustering analysis (**Supplementary Figure 8**).

Next, to identify tissue-specific AS events, we derived a AS event tissue-specificity score (AS event TS_Score) using tissue PSI/PSU/PIR values for each AS events in each tissue (See Methods for details), also taking into account the overall similarities between tissues. In total, 6445 AS events (6.7% of all AS events) in 4177 genes are tissue-specific (**Supplementary Table 5**), defined as having AS event TS_Score>2 (for tissue-specific inclusion events) or AS event TS_Score<−2 (for tissue-specific exclusion events) in at a subset of tissues. Biologically, AS event TS_Score>2 or <−2 can be interpreted as having the intron (in the case of intron retention) or exon at least 4 (i.e. 2^2^) fold more included or excluded in the target tissue than the other tissues on average. Statistically, this threshold represents the AS events of the top 1% TS_Scores. **Supplementary Figure 9** provides a heatmap overview of the PSI/PSU/PIR values of these tissue-specific AS events. Interestingly, unlike the tissue-specific gene expression observed in a large number of tissue types, tissue-specific splicing is mostly observed in a handful of tissues including testis, brain subregions, heart, skeleton muscle, whole blood, these tissue types together accounting for 82% of the tissue-specific AS events identified (Figure 6C). The rest of the tissue types show much fewer genes with tissue-specific AS events. This biased distribution of tissue-specific AS event is consistently observed even when we vary the stringency of AS event TS_Score thresholds (**Supplementary Figure 10**). Furthermore, the majority of tissue-specific AS events appeared to be tissue-specific AltEx events (27.5%) and IR events (67%), and very few are tissue-specific Alt5_3 events (4.5%).

A survey of targets with drugs in experimental and different clinical stages showed 72% of drug targets (**Supplementary Table 7**) have at least one AS event observed in the tissues examined, further confirming the importance of evaluating the tissue distribution patterns of alternative splicing for drug targets. There are numerous examples of alternative splicing affecting the pharmacology of the drug targets. For example, AMPA receptors are ionotropic transmembrane receptors for glutamate that play important roles in synaptic signal transmission. Drugs targeting AMPA receptors are being tested and used to treat a range of neurological diseases such as epilepsy and cognitive disorders ^30,31^. AMPA receptors consist of heterotetramers of four types of subunits (GRIA1-4). Alternative exon usage at the ‘flip-flop’ exons has been reported to have profound effect on receptor function, membrane trafficking, and receptor affinity to small molecule drugs ^32^. In the GTEx datasets, the alternative splicing between the ‘flip-flop’ exons can be clearly observed in all four AMPAR subunit genes in brain, but the spatial distribution of the two isoforms in the different brain subregions widely vary (**Supplementary Figure 11**). The ‘flop’ isoforms of GRIA2-4 are primarily expressed in cortex, caudate, and putamen (mean PSI *Ψ*=57%), while the ‘flip’ isoforms dominate in amygdala, hypothalamus, hippocampus, substantia nigra, and spinal cord (mean PSI *Ψ*=82%). Cerebellum has a mixed expression of the ‘flip’ isoforms for GRIA1,3 and the ‘flop’ isoforms for GRIA2,4. Thus, drugs preferentially targeting one of the isoforms might have profoundly different pharmacological effects on the different neurocircuits.

We are also particularly interested in leveraging the tissue-specific alternative splicing in therapeutic targets to uncover novel tissue specific targeting opportunities. A cross-examination of gene-level tissue specificity score and AS event tissue specificity score shows 1359 out of the 6445 (21%) tissue-specific AS events occur in the tissues where the genes are specifically expressed (gene TS_Score>3). The majority (79%) of tissue-specific AS events occur in tissues where genes not specifically expressed (Figure 6D), suggesting tissue-specific splicing could add a new dimension to tissue specific targeting. Here, we use CACNA1C gene as an interesting example of potentially leveraging tissue-specific splicing for tissue targeting. CACNA1C is subunit of L-type voltage-gated calcium channel (Cav1.2) responsible for the contractility of cardiomyocyte and vascular smooth muscles. Calcium channel blocker drugs targeting CACNA1C have been commonly used to treat hypertension. In brain, Cav1.2 is believed to regulate long-term potentiation (LTP). Recent human genetics studies have further implicated CACNA1C in psychiatric disorders such as schizophrenia and bipolar ^25,26,33^. Thus, there are strong interests to be able to specifically targeting CACNA1C in a brain in order to minimize potential cardiovascular safety liabilities. According to GTEx transcriptome altas, CACNA1C is highly expressed in all heart, artery, and brain tissues but not enriched in brain comparing to peripheral tissues (average gene TS_Score in brain regions is merely 0.02). However, a search for brain-specific AS event using our AS event TS_Score identified two AS events (exon8/8a, exon21/exon22) with varying degree of brain specificity. Among them, exon 21 and 22 are mainly mutually exclusive of each other where exon22-spliced in isoform is predominately expressed in brain (median PSI *Ψ*=91%, AS event TS_Score=4.26±0.11 in brain subregions). Based on the transmembrane structure prediction ^34^, exon22 is believed to encode a section of the extracellular loop and the adjacent transmembrane domain and significantly different from exon21 in seven amino acid residues. In addition, this structural difference between the two CACNA1C isoforms may also affect the receptor voltage-dependent kinetics as structure-activity effect caused by mutations has been reported in other L-type calcium channels ^35^. While it remains to be proven these structural differences could provide sufficient therapeutic window for brain-specific targeting, this exemplifies the potential utility of alternative splicing to derive for novel and testable hypothesis for tissue-specific targeting.

## Discussion

Understanding the tissue distribution of therapeutic targets and their splice isoforms is fundamentally important to every drug discovery program. The quantitative and comprehensive analysis of the tissue specificity of human gene expression and alternative splicing enabled by the GTEx transcriptome atlas is a major contribution to drug discovery and research community.

The GTEx transcriptome atlas is unprecedented in scale and quality, with deep RNA-seq profiling data generated on thousands of human tissue samples across a wide-range tissue types. Our analysis using a weighted tissue-specificity scoring metrics shows the GTEx transcriptome atlas vastly expanded the catalog of tissue-specific genes, and identified a large number of lowly expressed tissue-specific genes previously underestimated by commonly used microarray-based atlas. The systematic capture and categorization of these lower abundance tissue-specific genes is particularly important for studying tissues such as brain with diverse cell type compositions, where cell-type specific genes present in only a small fraction of cell populations are further diluted in expression and appear as low abundance at the tissue-level. From drug target discovery’s perspective, these low abundance tissue-specific genes could be particularly interesting. They are functionally important as indicated by our functional enrichment analysis, but may require much lower drug exposure to achieve full target occupancy and modulation, therefore, lowering the chance of off-target safety liabilities.

With the use of next-generation sequencing technology, deep sequencing coverage, and large sample size per tissue type, GTEx transcriptome atlas also enables a new dimension of tissue specificity analysis on alternative splicing. Our AS event-based splicing analysis of GTEx samples confirms previous observations that splicing is a universal phenomenon across human tissues. Examples like the differential splicing patterns in AMPA receptor subunits across brain tissues further highlight the importance of alternative splicing analysis of drug targets especially at the early stage of a drug discovery program.

Our joint analysis of gene-based and junction-based tissue-specificity scores (Figure 6D) indicates a large number of tissue-specific splicing occurs in genes that are not specifically expressed in the same tissues. This further suggests that both would offer complementary opportunities for tissue targeting. Particularly, the large number of tissue-specific splicing identified in brain, skeleton muscle, heart, and immune cells, could open up many more exciting new opportunities like the CACNA1C example for tissue-specific targeting. The unique tissue-specific splicing patterns in these tissues also suggest distinct regulatory mechanisms on alternative splicing, and the contribution of dys-regulated splicing to diseases relevant to these tissue types.

Tissue-specific gene expression has been instrumental in providing clues to understanding the molecular mechanisms underlying normal biological processes and human diseases. Here, by empirically examining the tissue expression pattern of drug targets, we further established tissue specificity as a desirable attribute for therapeutic targets. It is worth noting that targets currently before PhIII displayed a significant drop off in enrichment for high tissue-specificity comparing to PhIII+ targets (Figure 3B). We speculate that the coincidental overlap with the expected attrition in drug development pipeline may call for more attentions to tissue specificity early on. One explanation for the benefit of choosing tissue-specificity targets is that it bodes strong pre-clinical confidence for avoiding potential undesirable toxicity concerns ^36^. In practical terms, the chance of finding validated drug targets would have been drastically increased based on tissue-specificity criteria alone comparing to random (OR=2.5, Pvalue=3E-24, **Supplementary Figure 4A,B**). Furthermore, we compared the enrichment of PhIII+ targets when selecting based on “is this gene expressed in the disease tissue” vs. “is this gene preferentially expressed in the disease tissue”. As a broader first approximation, we assumed that the intended tissue for the target mechanism of action (MoA) is represented by the tissue with the highest TS_score for each drug target. The criteria to enforce tissue-specificity has a drastic improvement in enriching for PhIII+ targets comparing to the one requiring mere target being expressed in the tissues (**Supplementary Figure 12**). To further confirm the finding, we focused on only CNS targets as their MoAs could be more confidently assigned to brain. We selected targets classified by SOC terms “psychiatric disorders” and “nervous system disorders” and subsequently removed “Multiple Sclerosis” targets for the emergent opinion that it is an immunological disease. As shown in **Supplementary Figure 12B**, requiring brain-specific expression results in further enrichment of validated CNS targets over brain-expressed genes

In addition, we postulate that another explanation for the observed enrichment towards tissue-specificity could be the therapeutic availabilities of wider range of targets, particularly those with low expression levels. Comparing PhIII+ targets to the background, we observed that even though the overall expression distributions mirror each other, lowly-expressed PhIII+ targets are often tissue-specific (96% comparing to only 61% in background, p=2.2E-29, **Supplementary Figure 13A,B**). Our findings suggest that consideration for tissue-specificity may allow for the accessibility to wider range of targets (regardless of expression levels), an important consideration for target prioritization particularly in a subset of diseases (**Supplementary Figure 13C**).

This expanded catalog of tissue-specific genes also opens up opportunities to discover novel tissue-specific therapeutic targets. Our study of brain striatum-specific genes provides a useful template for mining human tissue specificity to identify potential therapeutic targets for human diseases. Drug development efforts often focused on a few gene families amenable to small molecule intervention (e.g., kinases, transporters, enzymes and receptors). Overlaying tissue-specificity can help further identify and prioritize disease-relevant targets among these druggable gene families. As an example, for central nervous system disorders, the large number of GPCRs identified in the striatum could help elucidate striatum-specific druggable targets and potential molecular regulators of this important brain region. Among them, several of the dopamine receptors identified are the molecular targets of approved medications including antipsychotics to treat psychiatric diseases and dopaminergic agonists to treat Parkinson’s disease. Genetic variations in the D2 receptor gene are also associated with schizophrenia risk in the recent meta-GWAS analyses ^25,26^. Interestingly, six of the striatal-specific GPCRs are orphan receptors whose functions are largely unknown; however, their highly specific expression pattern provides new clues suggesting these orphans might regulate striatal neurotransmission and function, and could also be attractive drug targets for psychiatric and Parkinson’s diseases. Indeed, one of the orphan receptors identified as GPR88 also showed conserved striatal expression in other vertebrates including rodents and non-human primates ^37^. Recent studies in mouse models indicate GPR88 negatively modulates the activity of striatal medium spiny neurons to affect memory and motor control ^38^ and small molecule agonists for GPR88 have been recently developed as potential therapeutics for diseases of the striatum/basal ganglia ^39^.

GWAS and other large-scale sequencing studies have elucidated the genetic architecture underpinning complex diseases. While this provides a plethora of unbiased causal connections between genetic variations and disease, gaps still exists in identifying the true causal genes and actual functional variants in each of these genetic risk loci. By seeking relationships between risk loci and tissue-specific gene catalog derived from the GTEx transcriptome atlas, we demonstrated that the enrichment of GWAS signals around disease-relevant tissue-specific genes is a strikingly common phenomenon among complex diseases, consistent with previous observations that GWAS variants tend to be enriched in tissue-specific regulatory elements ^40,41^. Thus, our enrichment analysis of tissue-specific gene expression in the 61 GWAS studies provides orthogonal evidence for prioritizing disease causal genes in GWAS loci.

Furthermore, the tissue-specific disease candidate genes identified through the GWAS-GTEx integrative analysis provides the basis for defining a cellular context for further functional studies. For example, the observation that blood and lymphocyte-specific genes are highly enriched in the GWAS loci of not only inflammatory diseases (e.g. IBD, Crohn’s disease, rheumatoid arthritis) but also diseases for other organs (e.g. Multiple sclerosis, T1D, Alzheimer’s diseases) further reinforces the causal contribution of immunity and inflammation as drivers of these complex diseases. Our findings corroborate well with the recent reports of overrepresentation of T cell–specific eQTLs among susceptibility alleles for multiple sclerosis and monocyte-specific eQTLs among Alzheimer’s disease variants^42^. Moreover, several blood-specific candidate genes in the Alzheimer GWAS loci (such as CD33, MS4A6A, INPP5D, CASS4) are specifically expressed at a high level in microglia, the brain resident immune cells ^43–46^. Overlap between genes specifically expressed in immune system and LOAD GWAS candidate genes supports the rationale to study their functional impact on microglia and mechanistically delineate the dys-regulated inflammatory pathways involved in the development of Alzheimer’s disease.

In closing, our genome-wide quantitative survey of tissue-specific gene expression using the GTEx transcriptome atlas provides a valuable resource for drug discovery applications. Integrative analysis anchored on tissue-specificity is an effective strategy for therapeutic target identification and evaluation, and enables further functional characterization of disease-causing genes.

## Methods

### Datasets

RNA-seq gene expression read counts and RPKM data were taken from GTEx consortium (http://www.gtexportal.org, v6p release). Detailed documentation of the sample collection, RNAseq data generation and processing, and gene expression quantification is available at the main consortium paper ^47^. A total of 8527 samples that passed QC filtering were used in the tissue-specificity analysis. Tissues with less than 10 samples were excluded. GENCODE v19 was used for gene types annotation and exon coordinates. Only protein-coding genes, long non-coding RNAs genes (lincRNA and antisense RNA) and pseudogenes (a total of 46508 genes) in the GENCODE v19 annotation were included in the gene expression TPM (Transcript Per Million) quantification and subsequent tissue-specificity calculation. We also assessed the impact of the inclusion of reads from highly abundant mitochondrial and globin genes on TS_Score calculation and downstream tissue-specificity analysis, as for a subset of tissues elevated levels of these abundant genes result in reduced gene expression values of other genes after TPM normalization. However, removing these reads from the calculations could also potentially remove factors biologically relevant to tissue-specificity assessment (e.g. coordination of mitochondria and nuclear gene expression). Here, we verified that the inclusion of these reads in our TS_Score calculations do not affect our main conclusions on the tissue-specific properties of drug targets and disease genes (**Supplementary Figure 14**). But as an additional resource, in **Supplementary Table 8**, we provided a version of TS_Scores where the read counts from mitochondria and globin genes were removed before the TPM and TS_Score calculations.

### Clustering of tissue types based on global gene expression patterns

First, gene read counts of each sample were normalized to TPM values (Transcript Per Million). For each gene, median TPM expression value was calculated for all samples in each tissue type to obtain the gene expression matrix *Exp_m,n_* (m genes, n tissues). Then the *Exp_m,n_* matrix is further log2 transformed and row median centered to obtain *Exp’_m,n_*, the relative gene expression value for each gene at each tissue. Pairwise Pearson correlation matrix between all tissues (*Cor_n,n_*) was calculated on *Exp’_m,n_* and then Ward’s hierarchical clustering was applied to *Cor_n,n_* to generate tissue groups using a tree-cutting threshold of 0.9.

### Tissue specificity calculation

For a given gene g, its tissue-specificity in tissue *t* (*TS_Score_g,t_*) is calculated according the following formula. 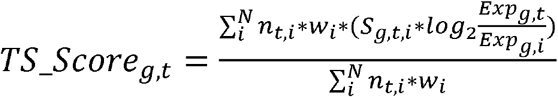 ***Exp_g,t_*** is the median TPM gene expression value for gene *g* in the target tissue *t* ***Exp_g,i_*** is the median TPM gene expression value for gene *g* in tissue *i*. ***log_2_(Exp_g,t_/Exp_g,I_*)** is the log_2_ ratio of the TPM gene expression values for gene *g* between target tissue *t* and tissue *i*, and is limited to [−5,5] range.

***w**_i_* is the weight for tissue *i* to adjust for the global gene expression similarity between tissue *i* and other tissues on the panel. To calculate this weight, pairwise Pearson correlation coefficients between tissues (*Cor_n,n_*) are first calculated using median centered gene expression matrix *Exp’_m,n_*. Then the weight for a given tissue *i* is calculated as one over the sum of the Pearson correlation coefficients between tissue *i* and other tissues i.e. *W_i_* = 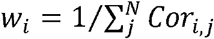. Correlation coefficients (*Cor_i,j_*) less than 0.4 are set to 0 so that only highly correlated tissues contribute to the weight.

***S_g,i_*** is a binary flag, set to 1 only when the log(*Exp_g,t_*/*Exp_g,i_*) value is statistically significant (FDR adjusted Pvalue<0.01). The significance is assessed using the linear model function ‘voom’ in the ‘limma’ package ^48^ designed for RNA-seq read counts and taking into account the variance among the samples.

***n_i_*** is also a binary flag to indicate whether tissue *i* is in tissue group different from the target tissue.(e.g. *n_i_*=0 when tissue *i* and tissue *t* are in the same tissue group). With the *n_i_* flag, tissues in the same group as the target tissue would not contribute to the tissue-specificity calculation for the target tissue.

To calculate brain sub-region specificity scores, the same specificity score formula was applied to only the 13 brain tissues. Tissue-specific gene was defined statistically as TS_score greater than 3, (>2 times standard deviation from the mean. ts_score μ=−0.05 and σ=1.28,) and can be biologically interpreted as having the expression in the target tissue 8 times of the weighted average expression of the rest of the tissues.

### Comparison with microarray-based transcriptome atlas from GNF

GNF data was downloaded from BioGPS. Gene expression values for the two array platforms (U133A, GNF1B) were combined. 25 tissue types on the GNF panel matched with the GTEx tissues based on their anatomical description. To calculate tissue-specificity score using the GNF panel, a modified formula was used with the *S_g,I_* always set to 1, as samples in GNF panel were pooled from multiple individuals, therefore, variance across samples is lost and the statistical significance of expression difference between tissues cannot be assessed. When comparing the GTEx and GNF datasets, tissue-specific genes were counted as unique in a dataset if they have TS_Score>3 in the dataset but TS_Score<1.5 in the other dataset.

### Functional annotation of tissue-specific genes

Gene Ontology functional enrichment in tissue-specific genes and genes with tissue specific splicing were carried out using Panther from GO Consortium ^49^. For enrichment of KO mouse phenotypes, KO mouse phenotype data was downloaded from MGI and genes with the same KO mouse phenotypes were group into the phenotype genesets. Then the enrichments of KO mouse phenotypes in the various tissue-specific genes were assessed using hypergeometric statistical tests in the R Package.

### Curation of drugs, targets, and disease ontology

Therapeutic program information was obtained from the⍰commercial Citeline Informa Pharmaprojects database (accessed March 2015). Programs with known human targets were parsed based on Entrez ID, which were then cross-referenced with Gencode 19 to ensure accuracy. Only unique targets are kept to exclude counting for statistics more than once. Clinical phases were assigned based on the furthest program across all documented indications. Disease indications from Citeline were mapped onto the Medical Dictionary for Regulatory Activities (MedDRA V17.1) Lowest-Level terms (LLTs), and output as the corresponding Primary System-Organ-Class (SOC) terms. The choice of PhIII+ targets (included “PhIII”, “Registered”, “Pre-registration”, “Launched”) for all statistical testing in this work ensured a conservative definition of “validated targets”. In the cases of comparisons by high-level ontology terms, choices were made to ensure that each target was assigned to only one disease term to avoid multiple-counting of the same target—the assignment tie-breaker was based on total number of PhIII+ drug programs followed by total number of all drug programs. In the cases of comparison of tissue-specificity scores, all genes (targets and background) were assigned the highest TS_Score across tissues.

### Enrichment of GWAS signals around tissue-specific genes

GWAS data were taken from NHGRI GWAS catalog Feb2014 version. In addition, recent meta-GWAS results for Alzheimer’s disease, Parkinson’s disease and schizophrenia were further added. Only GWAS lead SNPs with pvalue<5E-8 were included in the downstream analysis. GWAS SNPs of the same diseases or traits were further pruned by linkage disequilibrium (LD<0.8, LD derived from the 1000 Genome Project) and genomic proximity (distance<100kb) to ensure that each lead SNP marks an independent locus. 61 GWAS phenotypes with at least 15 genome-wide significant loci were selected to represent a diverse set of human diseases and traits. Genes located closest to the GWAS lead SNPs were identified based on the coordinates of the gene exons. For each human disease and traits, statistical significance of the overlap with genes closest to the GWAS lead SNPs against genes specifically expressed in each tissue type (TS_Score>3) was assessed using hypergeometric statistical tests.

### Alternative splicing event quantification and tissue-specific score calculation

We comprehensively quantified all major types of AS events involving alternative exons and exon skipping (AltEx), alternative 5’ and 3’ splice site selection (Alt5_3), and intron retention (IR) as previously described ^28,29,50^ and implemented in VAST-TOOLS (https://github.com/vastgroup/vast-tools). Briefly, we used the ‘align’ module from VAST-TOOLS to align the RNA-seq reads obtained from GTEx tissue samples to a comprehensive set of predefined splice junction library that include both curated spliced junctions from EST, cDNA, RNAseq transcripts and all hypothetically possible exon-exon junction combinations from annotated and de novo splice sites ^50^. Then for each sample, VAST-TOOLS identifies all AS events and quantifies them according to their types using the following metrics PSI: percent spliced in (for AltEx events), PSU: percent splice site usage (for Alt5_3 events), PIR: percent intron retention (IR events). PSI/PSU/PIR were calculated using exon-exon junction and exon-intron junction read counts corrected for sequence mappability at the junctions ^28^. **Supplementary Figure 7** shows schematic overview of the different types of AS events and the formula for the splicing metrics calculation. Each AS event is uniquely identified by the genomic coordinates of the exon-exon junctions and exon-intron junction involved. For each sample, AS events with insufficient read coverage using criteria defined in Irimia *et al*, 2014 were marked as non-quantifiable. Then, to quantify AS event in each tissue type, for each AS event, we calculated the tissue PSI/PSU/PIR value as the median PSI/PSU/PIR value across the samples with sufficient read coverage in each tissue type. AS events quantifiable in fewer than 5 samples of a tissue type were marked as non-quantifiable in that tissue. The total set of AS events across all tissues are defined as having 0.1< tissue PSI/PSU/PIR<0.9 in any of the tissues and/or having the range of tissue PSI/PSU/PIR values >0.25. And genes with multiple alternative spliced isoforms in individual tissue were defined as having at least one AS event with 0.1< tissue PSI/PSU/PIR<0.9 in the tissue.

Tissue specificity score (AS event ts_score) for each AS event is then calculated using the following formula, similar to the gene-level ts_score calculation, taking into account the global similarity between tissue types.

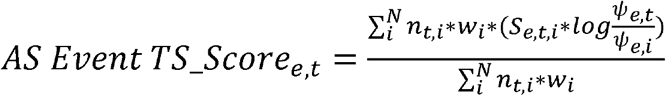

***ψ_e,t_*** is the tissue PSI/PSU/PIR value for AS event *e* at target tissue *t. A pseudovalue of 0.02 is added*.

***ψ_e,i_*** is the tissue PSI/PSU/PIR value for AS event *e* at tissue *I. A pseudovalue of 0.02 is added*.

***W**_i_* is the weight for tissue *i* to adjust for the global similarity of splicing patterns between tissue *i* and other tissues on the panel. Pairwise Pearson correlation coefficients (*Cor_i,j_*)of PSI/PSU/PIR values between tissues are first calculated. Correlation coefficients (*Cor_i,j_*) less than 0.4 are set to 0 so that only highly correlated tissues contribute to the weight. Then the weight for a given tissue *i* (*w_i_*) is calculated as one over the sum of the Pearson correlation coefficients between tissue *i* and other tissues i.e. 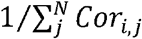.

***S_e,t,i_*** is a binary flag, set to 1 only when PSI/PSU/PIR values for AS event *e* in samples of target tissue *t* is statistically significantly different from samples of tissue *i* (FDR adjusted Pvalue<0.01), as assessed using the empirical Bayes log-odds of differential PSI/PSU/PIRs (^51^) (as implemented in “ebayes,” from the limma package in R), i.e. ***ψ_e,t_*** significantly different from ***ψ_e,i_***. Thus, this flag incorporates statistical significance assessment of the PSI/PSU/PIR value differences between tissues into the TS_Score calculation.

***N_t,i_*** is a binary flag, set to 0 if tissue *i* is in same tissue group as the target tissue *t*. (e.g. *n_i_*=0 when tissue *i* and tissue *t* are in the same tissue group). With the *n_t,i_* flag, tissues in the same group as the target tissue would not be compared to the target tissue in the tissue-specificity calculation. For example, a brain tissue would not be compared to other highly similar brain tissues.

***N*** is the number of tissues that AS event *e* is quantifiable.

Tissue-specific AS events are fined as having AS event TS_Score >2 or <−2, corresponding to top and bottom 1% of the TS_Scores and can be biologically interpreted as the PSI/PSU/PIR value in the target tissue being 4 (2^2^) folds of the average of other tissues. Since the PSI/PSU/PIR metrics are anchored on exon/intron spliced in, AS events with TS_Score<−2 are also defined as tissue-specific, as they represent cases where exon skipping or intron exclusion are tissue-specific. While the choice of TS_Score threshold stringency could vary, we further verified that the use of less stringent thresholds does not change the distribution pattern of tissue-specific AS events across tissues (**Supplementary Figure 10**).

## Acknowledgements

We would like to thank Robert Stanton, Enoch Huang, Christian Schubert, Jens Wendland, Jeff Struewing for their helpful suggestion to the manuscript. The Genotype-Tissue Expression (GTEx) Project was supported by the Common Fund of the Office of the Director of the National Institutes of Health. Additional funds were provided by the NCI, NHGRI, NHLBI, NIDA, NIMH, and NINDS. Donors were enrolled at Biospecimen Source Sites funded by NCI\SAIC-Frederick, Inc. (SAIC-F) subcontracts to the National Disease Research Interchange (10XS170), Roswell Park Cancer Institute (10XS171), and Science Care, Inc. (X10S172). The Laboratory, Data Analysis, and Coordinating Center (LDACC) was funded through a contract (HHSN268201000029C) to The Broad Institute, Inc. Biorepository operations were funded through an SAIC-F subcontract to Van Andel Institute (10ST1035). Additional data repository and project management were provided by SAIC-F (HHSN261200800001E). The Brain Bank was supported by a supplements to University of Miami grants DA006227 & DA033684 and to contract N01MH000028. Statistical Methods development grants were made to the University of Geneva (MH090941 & MH101814), the University of Chicago (MH090951, MH090937, MH101820, MH101825), the University of North Carolina - Chapel Hill (MH090936 & MH101819), Harvard University (MH090948), Stanford University (MH101782), Washington University St Louis (MH101810), and the University of Pennsylvania (MH101822). The data used for the analyses described in this manuscript were obtained from dbGaP accession number phs000424.v6.p1 on 01/20/2015.

## Author contributions

RY, JQ, HX conceived and designed the study. GTEx consortium generated the data. RY, JQ, RS, HX analyzed the data. RY, JA, HX wrote the paper. All main authors contributed to discussion, and helped revise the paper.

## References

1. Odom, D. T. et al. Control of pancreas and liver gene expression by HNF transcription factors. Science 303, 1378–1381 (2004).

2. Hwang, D. M. et al. A genome-based resource for molecular cardiovascular medicine: toward a compendium of cardiovascular genes. Circulation 96, 4146–4203 (1997).

3. Painter, M., Davis, S., Hardy, R. & Mathis, D. Transcriptomes of the B and T lineages compared by multiplatform microarray profiling. The Journal of... (2011).

4. Gautier, E., Shay, T., Miller, J., Greter, M. & Jakubzick, C. Gene-expression profiles and transcriptional regulatory pathways that underlie the identity and diversity of mouse tissue macrophages. Nature (2012).

5. ENCODE Project Consortium. An integrated encyclopedia of DNA elements in the human genome. Nature 489, 57–74 (2012).

6. Andersson, R. et al. An atlas of active enhancers across human cell types and tissues. Nature 507, 455–461 (2014).

7. FANTOM Consortium and the RIKEN PMI and CLST (DGT) et al. A promoter-level mammalian expression atlas. Nature 507, 462–470 (2014).

8. Thurman, R. E. et al. The accessible chromatin landscape of the human genome. Nature 489, 75–82 (2012).

9. Lage, K. et al. A large-scale analysis of tissue-specific pathology and gene expression of human disease genes and complexes. Proc. Natl. Acad. Sci. U.S.A. 105, 20870–20875 (2008).

10. Winter, D. R., Song, L., Mukherjee, S., Furey, T. S. & Crawford, G. E. DNase-seq predicts regions of rotational nucleosome stability across diverse human cell types. Genome Res. 23, 1118–1129 (2013).

11. Su, A. I. et al. A gene atlas of the mouse and human protein-encoding transcriptomes. Proc. Natl. Acad. Sci. U.S.A. 101, 6062–6067 (2004).

12. Wang, Z., Gerstein, M. & Snyder, M. RNA-Seq: a revolutionary tool for transcriptomics. Nat. Rev. Genet. 10, 57–63 (2009).

13. Mortazavi, A., Williams, B., McCue, K. & Schaeffer, L. Mapping and quantifying mammalian transcriptomes by RNA-Seq. Nature (2008).

14. GTEx Consortium. The Genotype-Tissue Expression (GTEx) project. Nat. Genet. 45, 580–585 (2013).

15. Kadota, K., Ye, J., Nakai, Y., Terada, T. & Shimizu, K. ROKU: a novel method for identification of tissue-specific genes. BMC Bioinformatics 7, 294 (2006).

16. Liang, S., Li, Y., Be, X., Howes, S. & Liu, W. Detecting and profiling tissue-selective genes. Physiol. Genomics 26, 158–162 (2006).

17. Schug, J. et al. Promoter features related to tissue specificity as measured by Shannon entropy. Genome Biol. 6, R33 (2005).

18. Santos, R. et al. A comprehensive map of molecular drug targets. Nat Rev Drug Discov 16, 19–34 (2017).

19. Lin, L., Yee, S. W., Kim, R. B. & Giacomini, K. M. SLC transporters as therapeutic targets: emerging opportunities. Nat Rev Drug Discov 14, 543–560 (2015).

20. Hopkins, A. & Groom, C. The druggable genome. Nat Rev Drug Discov (2002).

21. Russ, A. & Lampel, S. The druggable genome: an update. Drug Discov. Today (2005).

22. Haber, S. N. & Knutson, B. The reward circuit: linking primate anatomy and human imaging. Neuropsychopharmacology 35, 4–26 (2010).

23. Shepherd, G. M. G. Corticostriatal connectivity and its role in disease. Nat. Rev. Neurosci. 14, 278–291 (2013).

24. Plenge, R. M., Scolnick, E. M. & Altshuler, D. Validating therapeutic targets through human genetics. Nat Rev Drug Discov 12, 581–594 (2013).

25. Schubert, C. R., Xi, H. S., Wendland, J. R. & O’Donnell, P. Translating human genetics into novel treatment targets for schizophrenia. Neuron 84, 537–541 (2014).

26. Schizophrenia Working Group of the Psychiatric Genomics Consortium. Biological insights from 108 schizophrenia-associated genetic loci. Nature 511, 421–427 (2014).

27. Katz, Y., Wang, E. T., Airoldi, E. M. & Burge, C. B. Analysis and design of RNA sequencing experiments for identifying isoform regulation. Nat. Methods 7, 1009–1015 (2010).

28. Irimia, M., Weatheritt, R., Ellis, J. & Parikshak, N. A highly conserved program of neuronal microexons is misregulated in autistic brains. Cell (2014).

29. Braunschweig, U., Barbosa-Morais, N. & Pan, Q. Widespread intron retention in mammals functionally tunes transcriptomes. Genome... (2014).

30. Partin, K. M. AMPA receptor potentiators: from drug design to cognitive enhancement. Curr Opin Pharmacol 20, 46–53 (2015).

31. Rogawski, M. A. Revisiting AMPA Receptors as an Antiepileptic Drug Target. Epilepsy Currents 11, 56–63 (2011).

32. Ozawa, S., Kamiya, H. & Tsuzuki, K. Glutamate receptors in the mammalian central nervous system. Progress in neurobiology (1998).

33. Cross-Disorder Group of the Psychiatric Genomics Consortium. Identification of risk loci with shared effects on five major psychiatric disorders: a genome-wide analysis. Lancet 381, 1371–1379 (2013).

34. Liao, P., Yong, T. F., Liang, M. C., Yue, D. T. & Soong, T. W. Splicing for alternative structures of Cav1.2 Ca2+ channels in cardiac and smooth muscles. Cardiovasc. Res. 68, 197–203 (2005).

35. Lipscombe, D., Andrade, A. & Allen, S. Alternative splicing: functional diversity among voltage-gated calcium channels and behavioral consequences. Biochimica et Biophysica Acta (BBA)-... (2013).

36. Gashaw, I., Ellinghaus, P., Sommer, A. & Asadullah, K. What makes a good drug target? Drug Discov. Today (2012).

37. Massart, R., Guilloux, J.-P., Mignon, V., Sokoloff, P. & Diaz, J. Striatal GPR88 expression is confined to the whole projection neuron population and is regulated by dopaminergic and glutamatergic afferents. Eur. J. Neurosci. 30, 397–414 (2009).

38. Quintana, A., Sanz, E., Wang, W. & Storey, G. Lack of GPR88 enhances medium spiny neuron activity and alters motor-and cue-dependent behaviors. Nature (2012).

39. Jin, C. et al. Synthesis, pharmacological characterization, and structure-activity relationship studies of small molecular agonists for the orphan GPR88 receptor. ACS Chem Neurosci 5, 576–587 (2014).

40. Ernst, J. et al. Mapping and analysis of chromatin state dynamics in nine human cell types. Nature 473, 43–49 (2011).

41. Maurano, M. T. et al. Systematic localization of common disease-associated variation in regulatory DNA. Science 337, 1190–1195 (2012).

42. Raj, T. et al. Polarization of the effects of autoimmune and neurodegenerative risk alleles in leukocytes. Science 344, 519–523 (2014).

43. Malik, M. et al. CD33 Alzheimer’s risk-altering polymorphism, CD33 expression, and exon 2 splicing. J. Neurosci. 33, 13320–13325 (2013).

44. Bradshaw, E. M. et al. CD33 Alzheimer’s disease locus: altered monocyte function and amyloid biology. Nat. Neurosci. 16, 848–850 (2013).

45. Griciuc, A. et al. Alzheimer’s disease risk gene CD33 inhibits microglial uptake of amyloid beta. Neuron 78, 631–643 (2013).

46. Hickman, S. E. et al. The microglial sensome revealed by direct RNA sequencing. Nat. Neurosci. 16, 1896–1905 (2013).

47. GTEx Consortium et al. Genetic effects on gene expression across human tissues. Nature 550, 204–213 (2017).

48. Law, C. W., Chen, Y., Shi, W. & Smyth, G. K. voom: Precision weights unlock linear model analysis tools for RNA-seq read counts. Genome Biol. 15, R29 (2014).

49. Mi, H., Poudel, S., Muruganujan, A., Casagrande, J. T. & Thomas, P. D. PANTHER version 10: expanded protein families and functions, and analysis tools. Nucleic Acids Res 44, D336–42 (2016).

50. Han, H. et al. MBNL proteins repress ES-cell-specific alternative splicing and reprogramming. Nature 498, 241–245 (2013).

51. Smyth, G. K. in Bioinformatics and Computational Biology Solutions Using R and Bioconductor 397–420 (Springer-Verlag, 2005). doi:10.1007/0-387-29362-0_23

